# Super-resolution microscopy reveals the three-dimensional organization of meiotic chromosome axes in intact *C. elegans* tissue

**DOI:** 10.1101/112599

**Authors:** Simone Köhler, Michal Wojcik, Ke Xu, Abby F. Dernburg

## Abstract

When cells enter meiosis, their chromosomes reorganize as linear arrays of chromatin loops anchored to a central axis. Meiotic chromosome axes form a platform for the assembly of the synaptonemal complex (SC), and play central roles in other meiotic processes, including homologous pairing, recombination, and chromosome segregation. However, little is known about the three-dimensional organization of components within the axes, which consist of cohesin complexes and additional meiosis-specific proteins. Here we investigate the molecular organization of meiotic chromosome axes in *C. elegans* through STORM and PALM superresolution imaging of intact germline tissue. By tagging one axis protein (HIM-3) with a photoconvertible fluorescent protein, we established a spatial reference for other components, which were localized using antibodies against epitope tags inserted by CRISPR/Cas9 genome editing. Using three-dimensional averaging, we determined the 3D-organization of all known components within synapsed chromosome axes to a precision of 2-5 nanometers. We find that meiosis-specific HORMA-domain proteins span a gap between cohesin complexes and the central region of the SC, consistent with their essential roles in SC assembly. Our data further suggest that the two different meiotic cohesin complexes are distinctly arranged within the axes: Cohesin complexes containing COH-3 or -4 kleisins form a central core in the central plane of the axes, whereas complexes containing REC-8 kleisin protrude above and below the plane defined by the SC. This splayed organization may help to explain the role of the chromosome axes in promoting inter-homolog repair of meiotic double strand breaks by inhibiting inter-sister repair.

## Introduction

During meiosis, chromosomes undergo dramatic remodeling to enable homolog pairing, recombination, and segregation. A hallmark of meiotic entry is the reorganization of meiotic chromosomes into linear arrays of chromatin loops anchored to a central axis. The mechanism of this remodeling is not understood, but it involves replacement of canonical cohesin complexes with variant complexes containing meiosis-specific subunits. In addition to cohesins, other meiosis-specific proteins are recruited to chromosome axes and are required for their roles in synapsis and meiotic regulation (1, 2). While axis components have been identified and their interactions analyzed in various model organisms, their physical organization is poorly understood. Here, we use STORM (STochastic Optical Reconstruction Microscopy) and PALM (Photo-Activated Localization Microscopy) super-resolution methods (3-6) to investigate the architecture of chromosome axes in meiotic nuclei from intact gonads of *C. elegans.*

Meiotic chromosome axes form an essential substrate for the assembly of the synaptonemal complex (SC), which bridges the axes of paired homologs. In addition to their structural roles in reorganizing meiotic chromosomes and templating SC formation, axis proteins play a central role in meiotic chromosome dynamics: Axis assembly is required for homolog recognition as well as homolog-specific synapsis (7, 8). Furthermore, chromosome axes are required for double strand break (DSB) formation (9, 10) and are thought to regulate the processing of DSBs as they undergo recombinational repair. Specifically, axis structure and/or activities recruited by axis proteins inhibit the use of the sister chromatid as a template for homologous recombination, thereby promoting inter-homolog repair, which is essential for chiasma formation and proper homolog segregation (7, 11-14). The axis also recruits components of the DNA damage response pathway, which likely regulate both the abundance of breaks and the choice of recombination pathways (15, 16).

In *C. elegans,* four meiosis-specific HORMA-domain proteins (HTP-1, HTP-2, HTP-3 and HIM-3) localize to the axis, where they play distinct roles (7, 9, 17-19). Biochemical, structural and genetic evidence has revealed that these HORMA-domain proteins form a hierarchical complex. HTP-3 recruits HTP-1, HTP-2, and HIM-3 through interactions of their respective HORMA-domains with cognate closure motifs in the C-terminal tail of HTP-3 (20). However, how HTP-3 is recruited to the axis, and how these meiotic HORMA-domain proteins interact with cohesins is still unclear. Axis association of some meiotic cohesins and HTP-3 domain proteins are partially interdependent, indicating that these components might interact (9, 18, 21). Studies in other organisms have shown that an interdependence between HORMA-domain proteins and cohesin complexes is conserved among metazoans (22, 23).

Axial elements were described in early electron micrographs as electron-dense regions flanking the ladder-like central region of the SC, which is transversely striated and spans ∼100 nm (Fig. 1A, top) (24). However, recent evidence has indicated that the central region proteins alone organize into structures with longitudinal electron-dense components (25). This indicates that the electron-dense elements that flank the SC may not correspond to assemblies of axis proteins, although some axis proteins appear to localize near these structures (26). Recent work has illuminated some structural details of axis proteins and their interactions (13, 26-28). However, the overall organization of the chromosome axis remains largely undetermined.

**Figure 1:**
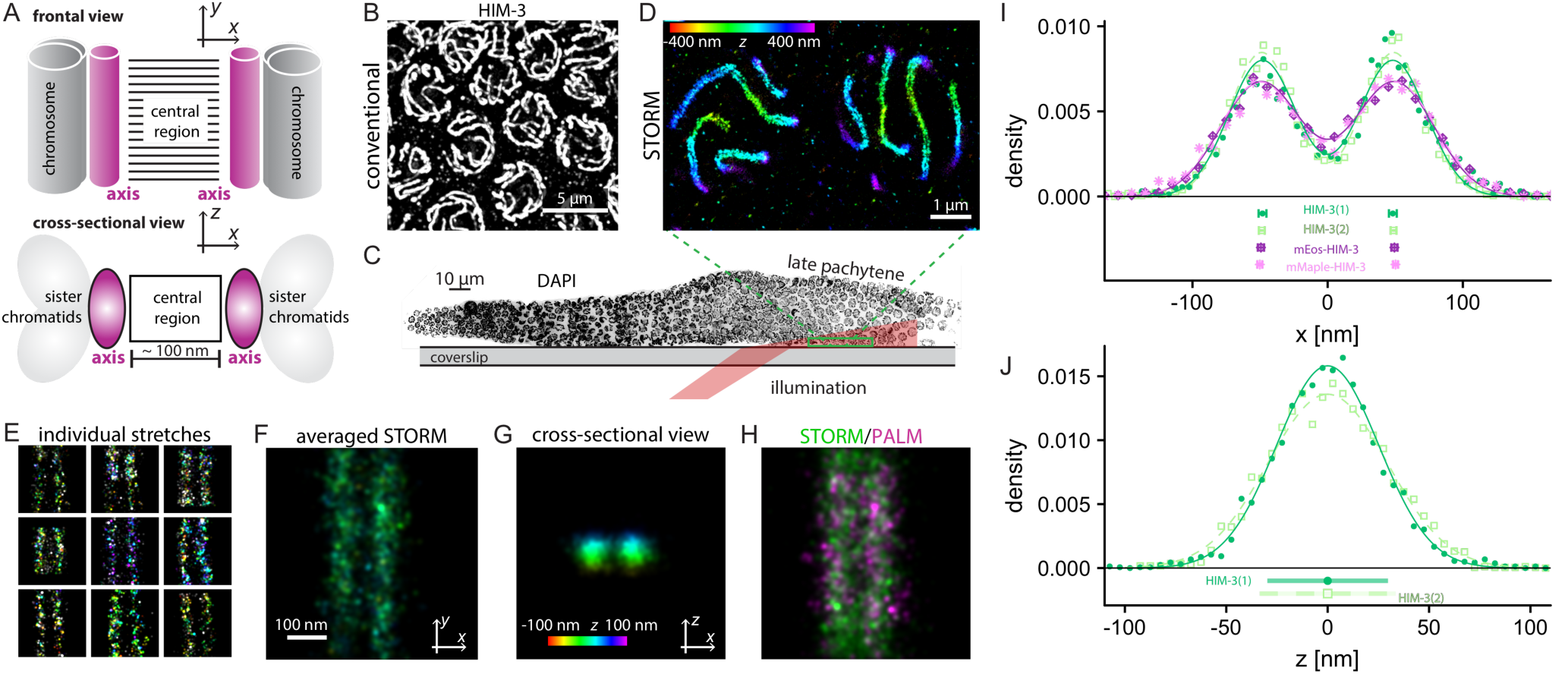
PALM and STORM localization microscopy of intact *C. elegans* gonads yields highly reproducible distance measurements. (A) In meiosis, chromosome axes (magenta) linearize homologous chromosomes (gray) and provide the platform for the assembly of the central region of the SC (black ladder). The cartoon depicts synapsed chromosomes in frontal *xy*-view (top), and in a crosssectional xz-view (bottom). (B) Conventional immunofluorescence microscopy cannot resolve paired chromosome axes stained for HIM-3, which are separated by about 100 nm. (C) Schematic illustrating how data were collected from *C. elegans* gonads. Tissue was pressed against a coverslip, and the nuclei closest to the coverslip were illuminated for STORM/PALM imaging. (D) Paired axes are readily resolved by 3D-STORM of HIM-3. Colors in 3D-STORM images indicate localization along the optical axis, with red being closest to the coverslip and violet at the most distant position (color scale bar). (E, F) To determine the localization of HIM-3, multiple individual stretches in frontal view were aligned to generate an averaged 3D-STORM image. Color denotes the position in *z* as shown by the color scale bar in (G). (G) Using optical astigmatism we determined the localization of HIM-3 in three dimensions, which can be visualized in a cross-sectional view of the chromosome axes. (H) The localization of HIM-3 determined by immunofluorescence with STORM (green) is indistinguishable from its localization based on PALM images of endogenously tagged mEos2-HIM-3 (magenta). (I) The distance between HIM-3 and the center of the SC was determined by fitting two normal Gaussian distributions to histograms of localization events in frontal view. HIM-3 is localized at 48.4±1.6 nm off center, which is highly reproducible (48.8±2.2 nm - dark green circles and 48.2±2.9 nm - light green squares, independent STORM experiments from two different days) and PALM experiments (mEos2: 49.4±2.3 nm - dark purple diamonds and mMaple3-HIM-3, HIM-1-intHA: 50.2±1.8 nm - magenta asterisks). Bars at the bottom of the graph represent the localizations of HIM-3 in different experiments with their respective standard deviations determined by bootstrapping. (J) The histogram of localization events in *z* reveals a well-defined localization pattern of HIM-3 across the chromosome axis, which is highly reproducible. The HWHM is 35±5 nm (dark green circles) and 30±3 nm (light green squares) in two independent experiments on different days. Bars at the bottom of the graph represent half widths at half maximum and their standard deviations estimated by bootstrapping.

Super-resolution microscopy has emerged as a powerful tool to bridge the resolving capabilities of conventional fluorescence and electron microscopy (29, 30). While the molecular organization of the chromosome axes cannot be resolved by diffraction-limited fluorescence methods, we have found that ultrastructural features of the axes can be probed by STORM/PALM super-resolution techniques. Because the architecture of meiotic chromosomes is highly regular and reproducible, we could combine these localization techniques with averaging methods (31-33) to attain a high-resolution molecular map of the chromosome axes to few-nanometer precision. Taking advantage of the well-characterized progression of meiosis within the germline of *C. elegans,* we determined the three-dimensional organization of synapsed axes in nuclei in intact germline tissue using a combination of STORM and PALM techniques. We use specific antibodies and CRISPR/Cas9-based genome editing to introduce epitope tags into genes encoding all known components of the chromosome axis, in some cases at multiple positions. We map the position of these epitopes with respect to an internal reference to reconstruct a detailed three-dimensional model of the synapsed chromosome axis. The structure that emerges from our data is fully consistent with known interactions among the constituent proteins. In particular, we find that meiosis-specific HORMA-domain proteins bridge the distance between cohesin complexes and the central region of the SC. Additionally, we detect intriguing differences between the conformation of distinct meiotic cohesin complexes, and provide evidence that these complexes show bilateral symmetry above and below the plane of each chromosome axis, suggesting how the axis may contribute to preventing recombination between sister chromatids.

## Results

### 3D-STORM in intact tissue

During the pachytene stage of meiotic prophase, homologous chromosome axes are held in parallel by the synaptonemal complex (SC), which assembles between them. The SC has a width of about 100 nm in most species and thus paired axes cannot usually be resolved by conventional fluorescence microscopy (Fig. 1B). While optical techniques such as structured illumination microscopy can resolve paired axes (34-37), higher resolution is required to determine the positions of individual proteins within the axes. We thus turned to super-resolution microscopy techniques based on the localization of single molecules (3-5, 38-40), which have resolved the positions of individual fluorophores at a resolution approaching 10 nm in plane (*xy*) and 20 nm in the axial (*z*) direction. This class of techniques was introduced as STORM (3) for dye-labeled samples and PALM (4, 5) for samples tagged by fluorescent proteins. Although STORM/PALM is more typically applied to thin samples, in this study we worked with intact *C. elegans* gonads extruded from adult animals (41). By adjusting the illumination angle to be slightly smaller than the critical angle of the glass-water interface (42, 43), we imaged several micrometers into the whole-mount samples. This enabled us to take advantage of the information about the meiotic stages of individual cells based on their position within this tissue, and to minimize potential artifacts arising from spreading or other sample preparation methods (44). Because paired axes are held at a precise distance from each other by the synaptonemal complex, we were able to localize proteins within these structures at significantly higher precision by analyzing the distributions of positions measured by STORM/PALM.

We first performed 3D-STORM on the axial HORMA-domain protein HIM-3, detected with an antibody raised against its C-terminus and labeled secondary antibodies conjugated to AlexaFluor 647. Intact gonads were extruded from adult hermaphrodites and sandwiched between a coverslip and a microscope slide in a small volume of imaging buffer, so that the tissue surface was apposed to the coverslip (see Methods). In most *C. elegans* meiocytes, the chromosomes are confined to a shell near the nuclear envelope, as the center of each nucleus is occupied by a large nucleolus. Thus, we were able to visualize chromosomes lying near the apical surface of these nuclei (closer to the basement membrane that surrounds the germline tissue). (Fig. 1C). We could readily resolve two parallel bands of HIM-3 staining separated by approximately 100 nm (Fig. 1D and Fig. S1). We found that in these images, synapsed chromosomes were predominantly oriented in frontal view, with the parallel axes lying at the same depth within the sample. This preferential orientation of paired chromosomes is likely due to their spatial confinement between the nucleolus and nuclear envelope. Nevertheless, some chromosome regions were oriented such that the plane of the SC was not parallel to the XY imaging plane. We excluded such regions from our analysis of protein organization within the axes, and used only regions observed in frontal view (*xy*-view, Fig. 1A top panel). Using optical astigmatism (6), individual proteins within these axial structures could be localized to a resolution of ∼20 nm in *xy* and ∼50 nm in *z*.

### Averaging over many samples increases the precision of super-resolution data

To determine the position of HIM-3 within paired, synapsed axes, we generated averaged images of localization events from multiple chromosome regions in frontal view (Fig. 1E,F). We fit the histograms of these localization events to two normalized, symmetrical Gaussian distributions (Fig. 1I). Using this approach (31-33) we determined the average position of components in the synapsed axes with a precision of a few nm. Based on this analysis, the center of each band of HIM-3 staining was localized to be 48.4±1.6 nm (standard deviation estimated by bootstrapping) from the midline of the SC, a value that showed remarkably little variance between samples prepared independently on different days (dark and light green lines in Fig. 1I). In electron micrographs, the central region of the SC in *C. elegans* spans about 96-97 nm (25, 45), Thus, the distance between the edge of this structure and it midline is 48 nm, and HIM-3 therefore localizes at or very close to this edge. This is consistent with evidence that HIM-3 is essential for SC assembly between chromosomes (13, 19), and suggests that it may directly interact with central region proteins.

Using optical astigmatism (6), we also determined the distribution of HIM-3 along the optical axis (*z*), i.e. perpendicular to the plane of the SC (Fig. 1A bottom panel). In these cross-sections (*xz* view) the distribution of HIM-3 in *z* within each axis was normal (Fig. 1G), with a half width at half maximum (HWHM) of 32±3 nm (Fig. 1J), which is comparable to its HWHM in *xy* of 29±1 nm. This indicates that HIM-3 is confined to a narrow plane in the center of the synapsed chromosome axes.

### Internal quality control by sequential STORM and PALM imaging

Because the orientation of parallel HIM-3 bands provides a robust, highly reproducible reference for the orientation of the SC, we used it as an internal standard in double-labeling experiments to define the organization of other axis proteins. Multicolor STORM in thick samples is challenging due to the high background and poor photoswitching performance of dyes outside of a narrow spectral regime. We therefore engineered photo-switchable fusion proteins to localize HIM-3 using mEos2 (46) and mMaple3 (47). The fluorescent tags were inserted at the 5’ end of the endogenous *him-3* coding sequence using CRISPR/Cas9 genome editing methods. The resulting N-terminal fusion proteins were fully functional, based on an absence of meiotic defects in worms expressing the fusion proteins in lieu of untagged HIM-3 (Table S1).

Using 3D-PALM, we determined the position of mEos2-HIM-3 to a resolution of 40 nm in *xy* (Fig. 1H,I). The separation of mEos2-HIM-3 and mMaple-3-HIM-3 from the SC midline were 49.4±2.3 nm and 50.2±1.8 nm (mMaple-3-HIM-3; HIM-1-intHA, see below), respectively (Fig. 1I), virtually identical to the distance determined by 3D-STORM using immunofluorescence. While most of our experiments were performed with mEos2-HIM-3, we note that mMaple3-HIM-3 has slightly superior photochemical properties. Moreover, its faster maturation time of about 30 min allowed imaging in earlier stages in meiotic prophase compared to mEos2, which restricted our observations to mid/late pachytene due to its maturation time of several hours (47). Nevertheless, the highly reproducible localization of HIM-3 across many samples and with both imaging methods provides evidence that the structure of the axes is not easily distorted during dissection, fixation, or mounting of intact gonad tissue for singlemolecule imaging. As the spatial resolution of PALM was lower than that of STORM, due to the limited photon yield from photoswitchable fluorescent proteins, we used STORM to analyze the localization of other axial components in whole gonads from worms expressing photoconvertible mEos2-HIM-3 or mMaple3-HIM-3, and used PALM images from the same field of view for structural alignment and quality control to assure constant localization of the reference protein HIM-3 in all experiments.

### Organization of meiosis-specific HORMA-domain proteins within the chromosome axis

HTP-3 is the largest of four meiotic HORMA-domain proteins expressed in *C. elegans.* It is recruited to the axis independently of HTP-1/2 and HIM-3 (9, 18, 20). HTP-1/2 and HIM-3 are recruited to the axis through binding of their HORMA domains to “closure motifs”, short peptide sequences within the long tail of HTP-3 (Fig. 2A). This direct physical interaction indicates that all four HORMA-domain proteins should be in close proximity within the chromosome axis. To test this hypothesis, we used STORM to map the positions of HTP-3 and HTP-1. Using a GFP antibody against a fully functional, C-terminally tagged HTP-3-GFP fusion protein (*htp-3(tm3655)* I; *ieSi6* II) (20) we detected two strands of HTP-3 at 54.7±2.1 nm off-center in frontal view, placing this epitope 6 nm farther from the center of the SC than HIM-3 (Fig. 2B top, E). Additionally, HTP-3-GFP was more broadly distributed along the z-axis, with a HWHM of 58±9 nm (Fig. 2B bottom, F). This broad distribution in *z* likely may reflect flexibility of the C-terminus of HTP-3, which is predicted to be largely unstructured. Interestingly, HIM-3, which binds to motifs in the middle of the long tail of HTP-3 (20), is constrained to the central plane of the axes. Given the known interaction of HIM-3 and the C-terminal tail of HTP-3, we wondered whether the distinct localization patterns of HIM-3 and HTP-3 might be caused by the insertion of epitope tags and/or use of antibodies. Alternatively, we wondered whether the long C-terminal tail of HTP-3, which spans almost 500 amino acids and has no predicted structured domains, might be specifically oriented within the chromosome axes. If so, we expected that the extreme N-terminus might be resolved from the position of the HIM-3-binding closure motifs in the center of its C-terminal tail. To test this hypothesis, we measured the position of the N-terminus of HTP-3 using a worm strain expressing 3xFlag-HTP-3-GFP inserted via MosSCI (48) *(htp-3(tm3655)* I; *ieSi62* II, Table S1). The N-terminus of 3xFlag-HTP-3-GFP was located at 60.4±1.8 nm from the SC midline in *x*, and in the center of chromosome axes, with a HWHM of 40±2 nm in *z* (Fig. 2C,E,F). Thus, HTP-3 is oriented diagonally within the chromosome axes relative to the plane of the SC, such that its C-terminus, which lies close to the central region, is widely distributed in *z*, while the HORMA-domain at its N-terminus is pointing away from the SC, and is confined to a narrow plane near the center of the SC (Fig. 2A).

**Figure 2:**
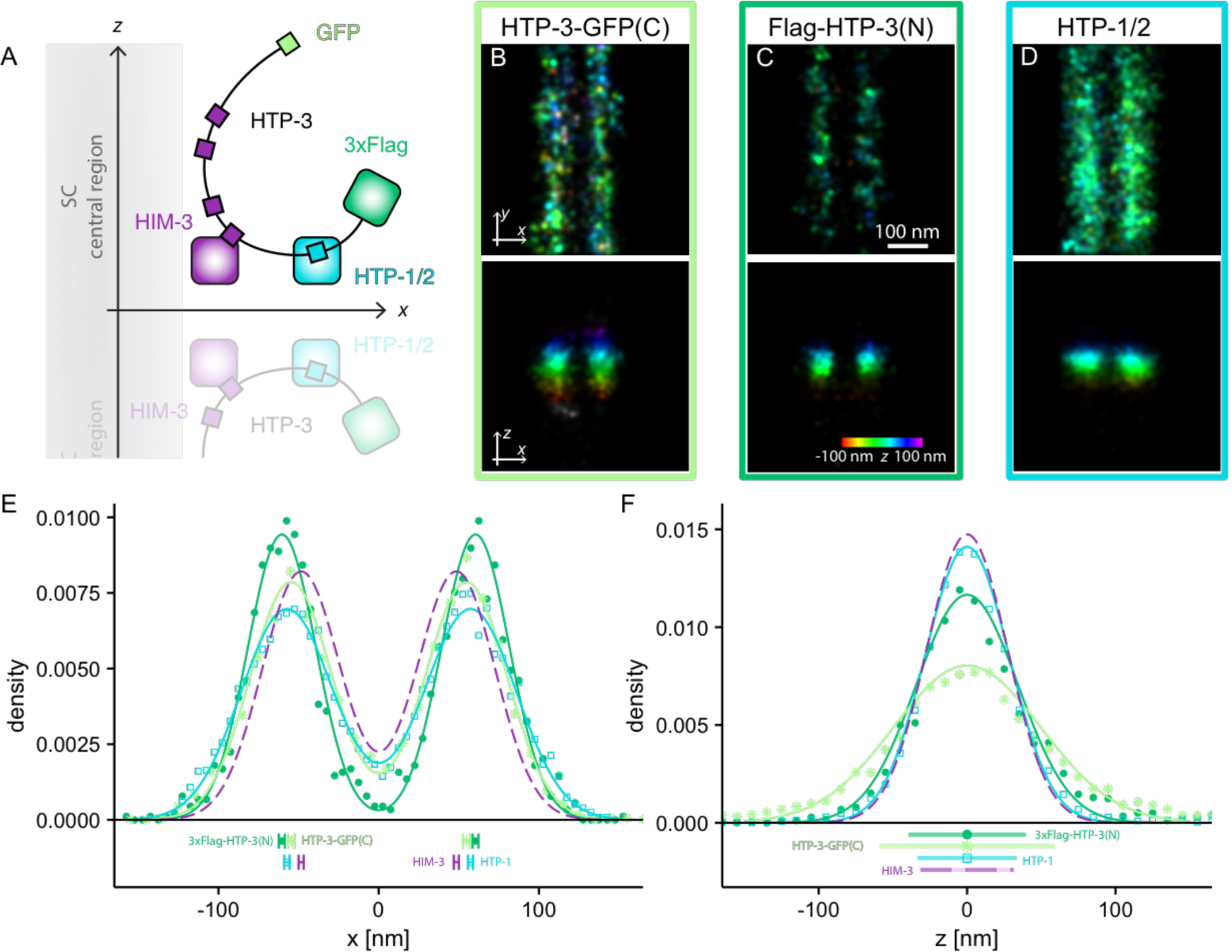
Localization of meiosis specific HORMA-domain proteins in the chromosome axes. (A) The positions of individual HORMA proteins within the axis, as determined by 3D-STORM, is consistent with the binding interactions among these proteins, as previously determined by *in vitro* and *in vivo* analysis (20). (B-F) The C-terminus of HTP-3-GFP (B, light green) localizes more proximally than the N-terminus (C, dark green), and shows a broader distribution in *z*, based on STORM images obtained in frontal view. HTP-1 (D, cyan) localizes closer to the C-terminus than the N-terminus of HTP-3, consistent with its binding to motifs in the C-terminal tail, but it shows a much narrower spatial distribution along the optical axis than a C-terminal GFP tag (top). (E, F) Distances were determined by fitting the distribution of localization events to double Gaussian distributions in *x* (E) and single Gaussians in *z* (F). (E) In frontal view, the distribution of HTP-1/2 (cyan squares) is centered at 57.3±1.6 nm from the SC midline, while the C-terminus of HTP-3-GFP (light green asterisks) lies at 54.7±2.1 nm off-center and the N-terminus of 3xFlag-HTP-3 (dark green solid circles) at 60.4±1.8 nm off-center. The position of HIM-3 (dashed purple line) is shown as a reference. Data points at the bottom of the graph indicate the localizations of HORMA-domain proteins with their respective standard deviations as determined by bootstrapping. (F) Across the axes in *z*, the HWHM is 33±2 nm for HTP-1/2 (cyan squares), 58±9 nm for HTP-3-GFP (light green asterisks) and 40±2 nm for the N-terminus of 3xFlag-HTP-3-GFP (dark green solid circles). HIM-3 (purple dashed line) is shown as a reference. The distributions are also represented as bars below the density profiles.

The HORMA-domain proteins HTP-1 and -2 are closely related paralogs arising from a recent gene duplication event. They share 83% sequence identity and are both recognized by a polyclonal antibody raised against HTP-1 (7). HTP-2 is not essential for proper execution of meiosis, but appears to overlap in function with HTP-1 (13). We therefore refer to this pair of paralogs as HTP-1/2. *In vitro* and *in vivo* experiments have demonstrated that HTP-1/2 are recruited to the chromosome axes through their interaction with closure motifs within the long tail of HTP-3 and at the extreme C-terminus of HIM-3 (20). Our aligned and averaged data confirm that HTP-1/2 localize to the chromosome axes at 57.3±1.6 nm off-center (Fig. 2D top, E). In frontal view, HTP-1/2 are thus between the N- and C-terminus of HTP-3, which is consistent with the interaction of HTP-1/2 HORMA-domain with closure motifs in the center of the tail of HTP-3. HTP-1/2 are narrowly distributed across the axes in cross-sectional view, with a HWHM of 33±2 nm, similar to both HIM-3 and the N-terminus of HTP-3 (Fig. 2D bottom, F). Thus, we observe distinct but similar localizations for each of the HORMA-domain proteins within synapsed chromosome axes, in both frontal and cross-sectional views, which are fully consistent with their direct physical interactions *in vitro* and *in vivo* (Fig. 2A).

### Organization of cohesins within the chromosome axis

While HORMA domain proteins play essential roles at chromosome axes in diverse phyla, they are not found in all clades. However, meiotic cohesins are required for axis function in all known species, and therefore likely govern their essential structure and functions, including the recruitment of HORMA domain proteins to the axis (9, 18, 49, 50). Therefore, we wondered whether STORM might illuminate how cohesin complexes are organized within the axes. Electron microscopy and crystallographic analysis have indicated that cohesins form a tripartite ring structure through the interactions of two large coiled-coil Structural Maintenance of Chromosome (SMC) proteins, known as SMC-1/HIM-1 and SMC-3 in *C. elegans* (51-54). Each of these proteins folds back on itself at its “hinge” domain to form an intramolecular superhelical coiled-coil, bringing together its N- and C-termini to form an ATPase domain, often called the “head”. The two SMC subunits form heterodimers through interactions at both their hinge and head domains. In addition, an essential “kleisin” subunit is thought to connect the head of one of the SMC proteins with the coiled-coil domain of the other (Fig. 3A). Proteolytic cleavage of the kleisin protein mediates release of cohesion during both mitosis and meiosis (55-57). In *C. elegans,* there are three meiosis-specific kleisins: REC-8, COH-3, and COH-4 (18, 21). COH-3 and COH-4 are highly similar and functionally interchangeable, but at least one of these proteins is required for proper axis function, as is REC-8.

**Figure 3:**
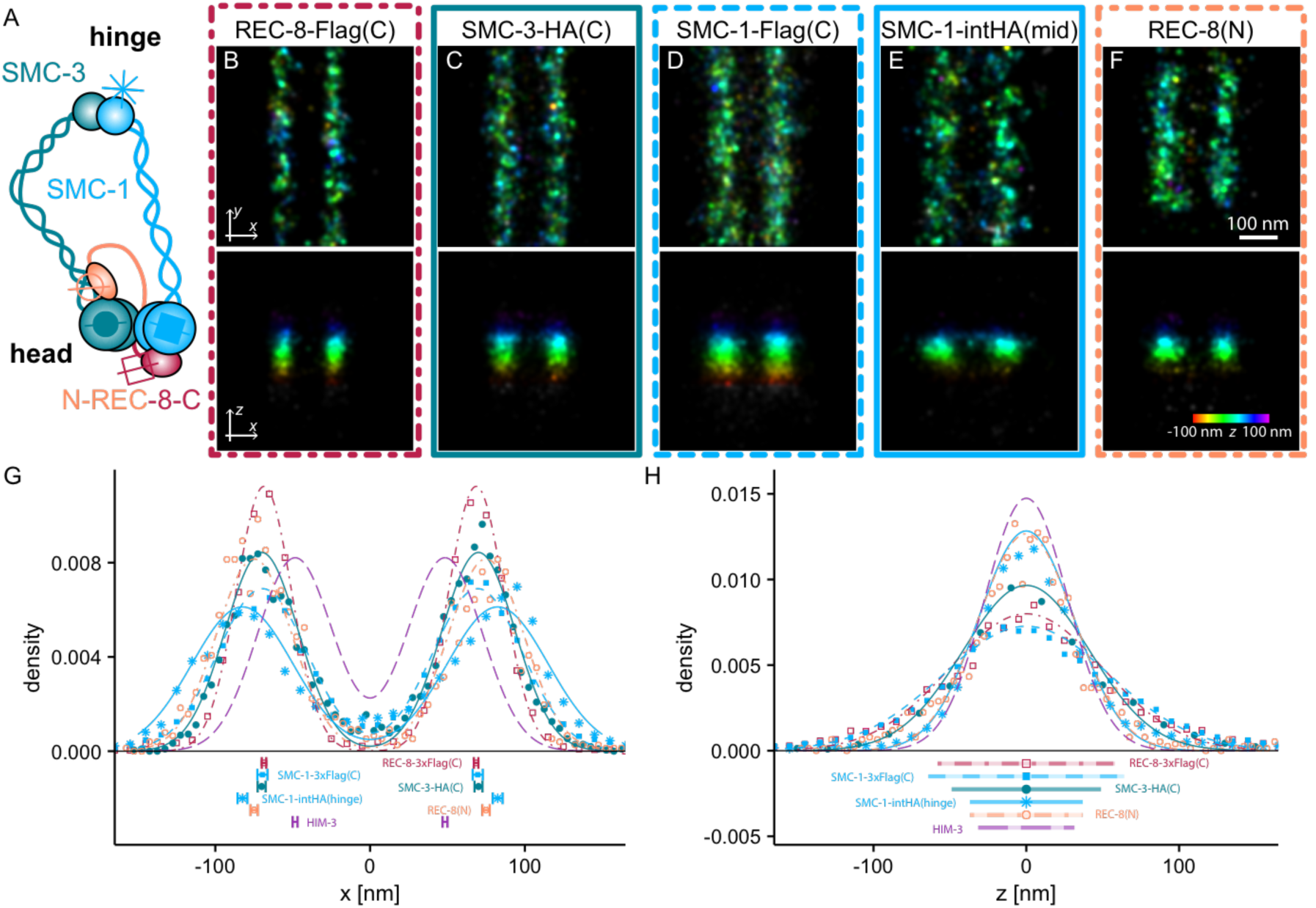
REC-8 cohesins have a defined orientation in the chromosome axes. (A) A schematic of the putative cohesin ring complex structure, as determined by single-particle EM imaging and crystallographic analysis, with the positions of the epitopes engineered in this study indicated. (B-F) STORM images of cohesin in the synapsed axes are used to measure the position of components in the axes in frontal (top) and cross-sectional (bottom) views. (G) Histograms of localization events in *x* show that all epitopes associated with the head domain localize to ∼ 70 nm from the SC midline, including the C-terminal REC-8-3xFlag tag (B, red open squares, 68.5±1.5 nm), C-terminal SMC-3-HA tag (C, teal closed circles, 70.1±2.6 nm) and C-terminal SMC-1-3xFlag tag (D, light blue closed squares, 69.6±3.2 nm). The hinge domain of SMC-1-intHA (E, light blue asterisks) has a mean distance of 82.5±3.2 nm and the N-terminus of the kleisin REC-8 (F, orange open spheres), which binds the coiled-coil domain of SMC-3, lies 74.9±2.4 nm from the midline. For reference, the localization of HIM-3 is shown (purple dashed line). (H) Epitopes associated with the cohesin head domain are broadly distributed along the optical axis (REC-8-3xFlag - 59±4 nm, SMC-3-HA - 49±7 nm, SMC-1-3xFlag - 65±5 nm) while SMC-1- intHA in the hinge domain and the N-terminus of REC-8 exhibit narrower HWHMs in *z* of 37±3 nm and 37±5 nm, respectively.

Crystal structures of the mitotic cohesin complex show that the SMC-1 and SMC-3 head domains and the C-terminus of the kleisin subunit form a tight complex (53, 58). Consistent with this, epitope tags that we introduced at the C-termini of REC-8 (REC-8-3xFlag), SMC-1 (SMC-1-3xFlag) and SMC-3 (SMC-3-HA) (Table S1) were localized in close proximity to each other at 68.5±1.5 nm, 69.6±3.2 nm and 70.1±2.6 nm off-center, respectively, in frontal view. They also showed similar distributions along the optical axis, with HWHM values of 59±4 nm, 65±4 nm and 49±7 nm, respectively (Fig. 3B-D,G,H). We therefore conclude that cohesin complexes in the synapsed chromosome axes are likely in a closed ring conformation, as predicted from the crystal structures. We next asked whether cohesin ring complexes, with an estimated diameter of 65 nm based on EM (51), have a specific orientation within synapsed chromosome axes. To this end we inserted an HA epitope tag into a poorly conserved loop in the hinge domain of SMC-1. This tagged protein supported the development and fertility of *C. elegans,* although we did observe a slight decrease in egg viability and an increase in male self-progeny from homozygous mutant animals, indicative of compromised function in mitosis and/or meiosis (Table S1). Nevertheless, the localization of HIM-3 was unaffected, suggesting a largely normal axis structure (Fig. 1I, mMaple-HIM-3). In frontal view, the hinge domain is localized 82.5±3.2 nm off-center (Fig. 3E top, G) and is thus separated by only 13 nm from the head domains. In cross-sectional view, the cohesin hinge domain is localized to a narrow plane in the center of the chromosome axes with a HWHM of only 37±3 nm (Fig. 3E bottom, H). In a crystal structure of the yeast cohesin complex (54), the N-terminal domain of the kleisin subunit REC-8 interacts with the coiled-coil domain of SMC-3. Thus, we expected that the N-terminus of REC-8 should be closer to the hinge domain than other components of the cohesin head (Fig. 3A). Indeed, the HWHM of localization events for an antibody recognizing aa 171-270 at the N-terminus of REC-8 in *z* is 37±5 nm. This is narrower than for other components in the cohesin head domain, and in frontal view it is located between head and hinge domain at 74.9±2.4 nm from the midline of the SC (Fig. 3F-H). This suggests that REC-8 cohesins are oriented diagonally within the chromosome axis, with their hinge domains more distal relative to the SC, and the head domains protruding towards the HORMA-domain proteins and the central region of the SC.

### Localization of COH-3/4 cohesin complexes

Meiotic kleisins play multiple roles, both during and prior to chromosome segregation. Orthologs of REC-8 were first identified in fungi, where they were found to hold sister chromatids together near their centromeres during the first meiotic division. Similarly, shortly before the first meiotic division in *C. elegans* oocytes, cohesin complexes containing REC-8 become asymmetrically enriched along the “long arm” of bivalents, where they hold sisters together during the first division (59). Conversely, complexes containing COH-3/4 become enriched along the “short arm,” and are targeted for cleavage to enable reductional segregation (disjunction of homologs) during the first division. Thus, specialized kleisins can mediate the unusual, stepwise release of cohesion required for proper partitioning of homologous chromosomes and sister chromatids to form haploid gametes. However, during early meiotic prophase, both types of cohesin complexes containing either REC-8 or COH-3/4 are distributed along the length of chromosomes. Deletion of *rec-8* has distinct effects on chromosome interactions from deletion of *coh-3/4,* indicating that even in early prophase, these cohesin complexes play different roles.

We therefore investigated whether REC-8 and COH-3/4 cohesin complexes are distinctly organized within the axes. In frontal view, the localization of a C-terminal HA tag on COH-3 (Table S1) was located indistinguishably from a C-terminal 3xFLAG epitope on REC-8, at 71.7±3.1 nm. However the COH-3 C-terminus showed a significantly narrower distribution along the optical axis, with a HWHM of 46±4 nm when compared to that of REC-8-3xFlag (59±4 nm) (Fig. 4A,D,E).

**Figure 4:**
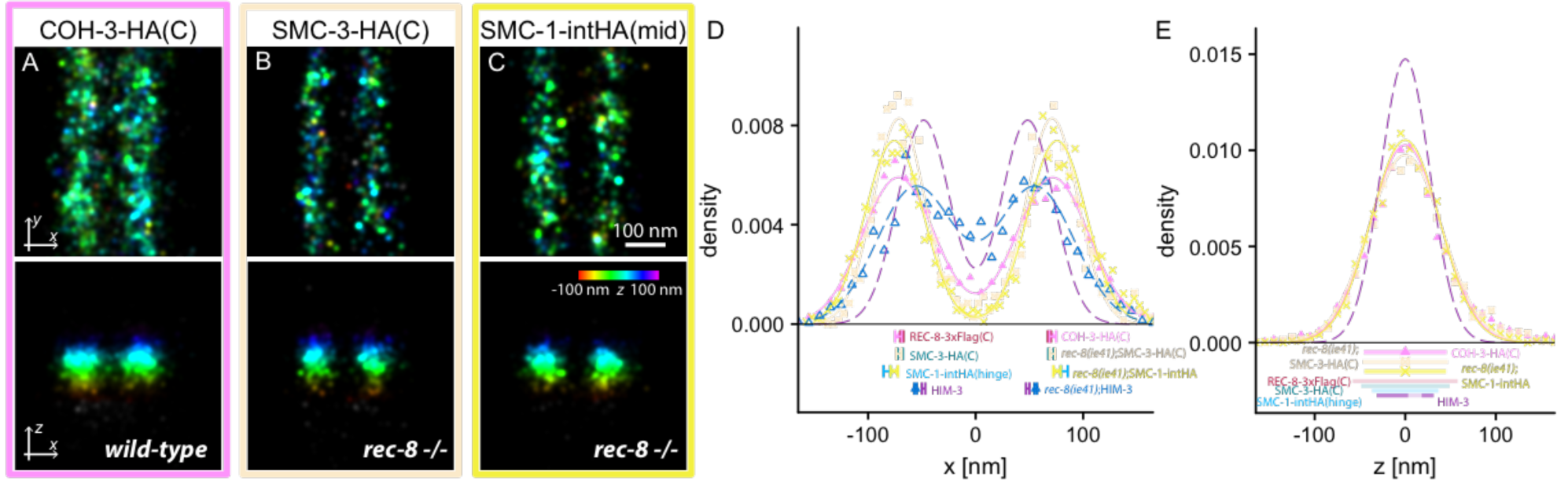
COH-3/4 cohesin complexes are localized distinctly from REC-8 cohesin complexes within chromosome axes. (A) In frontal view (top), the C-terminus of the kleisin COH-3-HA co-localizes with the C-terminus of REC-8 while its vertical distribution (bottom) is significantly narrower. (B) Similarly, the HWHM of SMC-3-HA is decreased in absence of REC-8, while the hinge domain of SMC-1-intHA (C) is shifted closer to the center in frontal view and slightly wider in *z* in *rec-8(ie41).* The positions in *x* and the HWHM in *z* are quantified in (D) and (E), respectively. The positions for COH-3-HA (pink solid triangles), *rec-8(ie41);SMC-3-HA* (tan crossed squares) and *rec-8(ie41);SMC-1-intHA* (yellow crosses) in *x* (D) are is at 71.7±3.1 nm, 71.2±2.6 nm and 75.7±3.4 nm, respectively, and their HWHM in *z* (E) are 46±4 nm, 48±1 nm and 45±3 nm. For reference, the positions and HWHM of REC-8-3xFlag (red), SMC-3-HA (teal) and SMC-1-intHA (light blue) in wild-type are shown in the summaries at the bottom of the graphs. HIM-3 in wild-type is shown in purple. Notably, mEos-HIM-3 is more distal in *rec-8(ie41)* in frontal view at 55.9±1.2 nm (navy open triangles).

It was not possible to analyze cohesin organization within chromosome axes in the absence of COH-3/4, since the chromosomes do not synapse when both genes are deleted, and the methods used here require a regular, symmetrical structure. However, we analyzed the organization of COH-3/4 cohesin complexes in the absence of REC-8, which was possible because *rec-8* mutants still undergo synapsis, although this does not appear to be fully normal, and may involve some inter-sister or non-homologous synapsis (18, 21, 59, 60) (see below). To this end, we used CRISPR/Cas9 mutagenesis to delete *rec-8* in both SMC-3-HA; mEos2-HIM-3 and SMC-1-intHA; mEos2-HIM-3 strain backgrounds. The mutation *rec-8(ie41)* introduces a stop codon and a frameshift mutation after the fourth amino acid in the REC-8 coding sequence, and should therefore recapitulate the phenotypes observed in other *rec-8* loss-of-function strains. Indeed, chromosomes synapsed in *rec-8(ie41)* (Fig. S2A-B). Interestingly, synapsis seems to be homologous in *rec-8(ie41),* at least for the X chromosome, since we detect a single, paired focus of HIM-8, which recognizes the pairing center of chromosome X (Fig. S2C-D). However, chromosomes in *rec-8(ie41)* mutant animals failed to form functional chiasmata, as indicated by the presence of univalents and chromosome fragments at diakinesis (Fig. S2E-F). Additionally, univalents at diakinesis showed a “butterfly”-like morphology, indicative of less extensive cohesion than in most other crossover-defective mutants, but identical to other *rec-8* mutant alleles (18, 21, 59). As suggested by conventional fluorescence images, super-resolution microscopy revealed a fairly normal organization of chromosome axes in *rec-8(ie41)* mutants (Fig. 4B-C). The localization of the head domain of SMC-3-HA (Fig. 4B) at 71.2±2.6 nm in frontal view (wild-type: 70.1±2.6 nm) and a HWHM of 48±1 nm in *z* remained largely unchanged in absence of REC-8. By contrast, the hinge domain showed a significantly wider distribution in *z*, with a HWHM of 45±3 nm (wild-type: 37±3 nm) (Fig. 4C) and was shifted closer to the head domain, at 75.7±3.4 nm off center in frontal view (wild-type: 82.5±3.2 nm). Moreover, the distribution of the hinge domain in frontal view showed a broader distribution in wild-type axes (Fig. 3G, HWHM in *x* of 38±3 nm) that was reduced in *rec-8(ie41)* (HWHM of 32±3 nm). A possible explanation for this difference is that the hinge domains of COH-3/4 and REC-8 cohesin complexes occupy different positions, and that the wide distribution seen in wild-type animals reflects a mixture of COH-3/4 and REC-8 cohesin complexes. In the absence of REC-8, both the hinge and head domains were more tightly distributed in *x* and *z*. This could indicate that COH-3 cohesin complexes form a central core along the length of the chromosome axis, while REC-8 mediates a splayed orientation of cohesin complexes within each axis, giving rise to a broader distribution along the optical axis.

Previous work has shown that the HORMA-domain protein HTP-3 is required for robust loading and/or stability of REC-8 cohesin complexes along the axis (18, 20). In our experiments, the position of the HORMA-domain protein mEos2-HIM-3 was shifted significantly farther from the SC in the absence of REC-8, from 49.4±2.3 nm to 55.8±1.2 nm off-center (Fig. 4D). Our results thus suggest an interplay between REC-8 cohesins and HORMA domain proteins, such that each affects the others’ organization and/or stability within the axis.

## Discussion

Using STORM/PALM we have mapped the localization of all known protein components of meiotic chromosome axes in *C. elegans,* in some cases with multiple epitope tags inserted at distinct positions (Fig. 5A). All epitopes on cohesin complexes localized more distally, relative to the central region of the SC, than epitopes on meiosis specific HORMA-domain proteins (Fig. 5B-C and supplemental Video S1). This key result indicates that the HORMA domain proteins bridge the distance between cohesins and the SC central region.

**Figure 5:**
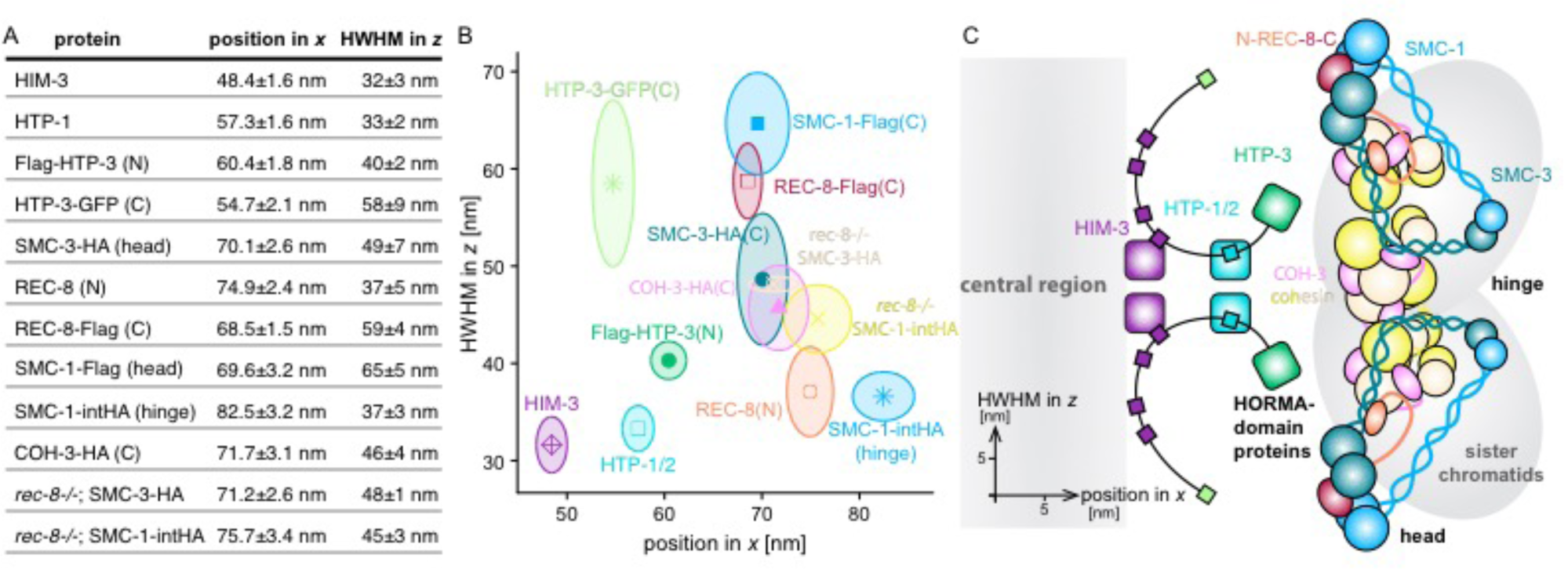
Model of synapsed chromosome axes in cross-sectional view. The positions from the midline of the SC in *x* and the HWHM in *z* of components within the chromosome axes measured by STORM are summarized in a table (A) and graphically (B). Mean values are shown by symbols and shaded areas are their corresponding standard deviations. These positions were used to construct a model of the synapsed chromosome axis (C); see the accompanying video for additional information about the mode

Overall, we find that the width of axial elements in frontal view spans at least 34.1 nm within the plane of the SC, the distance from our most proximal marker, HIM-3, to the most distal epitope on the cohesin complex. Epitopes that mark the four meiosis-specific HORMA-domain proteins, which form a flexible complex (20), span at least 12 nm in *x*, and cohesin complexes, which span 14 nm, are more distal from the central region and are clearly separated from the HORMA-domain proteins. The localization patterns we observe in synapsed chromosome axes *in vivo* are fully consistent with all genetic and biochemical evidence for interaction of components in the axis: The N-terminus of HTP-3, and thus its HORMA-domain, is in proximity to cohesin proteins, suggestive of a direct interaction, which is also supported by immunoprecipitation experiments (20) (Fig. 5B-C and supplemental Video S1).

HIM-3 and HTP-3 are strictly required for the assembly of the central region of the SC between chromosomes (7, 9, 17-19, 61). Our localization data demonstrate that the C-terminus of HTP-3, as well as the HORMA domain of HIM-3, are near this interface and might thus form a molecular platform for the assembly of the central region (Fig. 5B-C). While HORMA-domain proteins remain strongly associated with synapsed chromosome axes in *C. elegans,* they become depleted from axes upon synapsis in budding yeast and mice (62, 63). However, these proteins do not completely disappear from synapsed axes, and may therefore provide a persistent bridge between cohesins and the SC central region also in other organisms. Consistent with this idea, cohesin complexes and proteins of the central region were found to be separated by about 20 nm in mouse spermatocytes (28) suggesting that HORMA-domain proteins might indeed form a conserved interface between central region proteins and cohesin complexes.

Through analysis by 3D-STORM/PALM, we find that the axes show evidence of bilateral symmetry along the optical axis. Our data indicate that REC-8-containing cohesin rings are diagonally oriented in the axes, extending from a distal hinge domain in the central plane of the axis to bilaterally protruding head domains more proximal to the HORMA-domain protein complexes (Fig. 5B-C and supplemental Video S1). Based on a variety of *in vitro* evidence and mutational analysis, cohesion has been explained by a simple model in which two DNA molecules are entrapped by individual cohesin rings (58, 64, 65). Other models posit a requirement for interactions among cohesin complexes (reviewed in (66-68)). Supporting the latter possibility, a recent study demonstrated interallelic complementation of mutations in cohesin subunits for mitotic function in budding yeast (69). Interactions between cohesin complexes have not been reported *in vitro*, perhaps because they depend on chromatin association. Intriguingly, the bilateral localization pattern we observe for REC-8 cohesin complexes with central hinge domains and protruding head domains is consistent with such a higher order organization of cohesins *in vivo* (Fig. 5C and supplemental Video S1).

By contrast to the bilateral symmetry and specific orientation of REC-8 cohesin complexes, hinge and head domains of COH-3/4 cohesin complexes are indistinguishable in *x* and *z* cross-sections of the axis in absence of REC-8, which is consistent with COH-3/4 cohesin complexes forming a central cohesin core which is embedded by REC-8 cohesin complexes, which are in a splayed orientation (Fig. 5C and supplemental Video S1). Notably, electron micrographs have suggested that chromosome axes in rat spermatocytes might have three distinct layers, with a central layer perhaps acting as an interface connecting axes of sister chromatids (70). This is consistent with our findings, and suggests that a three-dimensional organization of chromosome axes with central and bilaterally protruding components is conserved across species.

In meiosis, chromosome axes play a central role in homologous recombination by shifting the bias of DSB break repair from inter-sister to inter-homolog recombination (11, 12, 61, 71, 72). Our finding suggesting that axes are bilaterally symmetric and are specifically organized in frontal view and across the axis can shed some light on the mechanism of this inter-homolog bias: cohesin complexes and HORMA-domain proteins may form a layer between sister-chromatids to prevent untimely sister-chromatid recombination while the establishment of a synapsis-competent surface by HORMA-domain proteins might promote inter-homolog recombination.

## Acknowledgements

Some nematode strains used in this work were provided by the Caenorhabditis Genetics Center, which is funded by the NIH - Office of Research Infrastructure Programs (P40 OD010440). This work was supported by a postdoctoral fellowship of the Human Frontier Science Program to SK (LT000903/2013- C), an NSF Graduate Research Fellowship (DGE 1106400) to MW, the Pew Biomedical Scholars Award and the Packard Fellowship for Science and Engineering to KX, and support to AFD from the National Institutes of Health (R01 GM065591) and the Howard Hughes Medical Institute.

## Author Contributions

SK and MW performed experiments and analyzed data. SK, MW, KX and AFD designed experiments and wrote the manuscript.

## Methods

### Worm strains and transgenes

All *C. elegans* strains were cultured using standard methods (73). Worms were maintained at 20°C. Strains used in this study are listed in Table S1. HTP-3-GFP (20) and 3xFlag-HTP-3-GFP transgenes were inserted by MosSCI (48) and crossed into mutants lacking an intact *htp-3* gene. For genome editing we initially performed CRISPR-Cas9 as described (74), which resulted in editing efficiencies of about 1% or less of F1 progeny positive for co-injection markers. Editing efficiencies were dramatically improved, to about 50% of marker-positive F1s, by injecting gRNA-Cas9 ribonucleoprotein (RNP) complexes pre-assembled *in vitro.* Cas9 protein was purchased from the UC Berkeley Macrolab, and Alt-R™ tracrRNA and crRNA were purchased from IDT. Equimolar solutions of tracrRNA and crRNA (100 μM each) were heated (95°C, 5 min) and annealed (room temperature, 5 min). Cas9-RNP was then formed by addition of purified Cas9 protein to a final concentration of 30 μM gRNA and 28 μM Cas9 protein for five minutes at room temperature. Repair templates were provided as plasmid (mEos2-HIM-3), long ssDNA (mMaple3-HIM-3 (25)) or ssODNs (synthesized as “Ultramers” by IDT) for epitope tagging and *rec-8(ie41),* respectively. All coding sequences for epitope tags and fluorescent proteins were codon-optimized for *C. elegans,* and the protein coding sequences also contained short introns (75).

### Immunofluorescence

Immunostaining of extruded gonads from adult hermaphrodites at 24 h post L4 was performed as previously described (41) with minor modifications. Worms were immobilized using 0.02 % tetramisole (instead of azide) during dissection. Fixed tissue was transferred to small polypropylene tubes, and all staining steps were carried out in suspension. Stained tissue was then mounted in ∼7 μL imaging buffer (Tris-HCl, pH 7.5, containing 100 mM cysteamine, 5% glucose, 0.8 mg mL^−1^ glucose oxidase, and 40 μg mL^−1^ catalase). The following primary antibodies were used: anti-HIM-3 (aa 154-253) (rabbit SDQ4713, 1:500, ModENCODE project (76)), anti-HTP-1/2 (rabbit, 1:500), anti-REC-8 (aa 171-270) (rabbit SDQ3914, 1:10000, ModENCODE project (76)), anti-GFP (mouse monoclonal, 1:500, Roche Cat#11814460001), anti-FLAG (mouse monoclonal M2, 1:500, Sigma) and anti-HA (mouse monoclonal 2-2.2.14, 1:500, ThermoFisher Scientific). Secondary antibodies for STORM are Alexa Fluor 647-anti-Rabbit (Goat or Donkey, 1:500, Invitrogen) or Alexa Fluor 647-anti-Mouse (Donkey, 1:500, Jackson ImmunoResearch).

#### Correlative STORM and PALM Imaging

3D STORM imaging was performed on a custom microscope based on a Nikon Eclipse Ti-U inverted optical microscope, using an oil immersion objective (Nikon CFI Plan Apochromat λ 100x, NA 1.45). Briefly, lasers at 647 nm (MPB Communications), 560 nm (MPB Communications), 488 nm (Coherent), and 405 nm (Coherent) were coupled into an optical fiber after an acousto-optic tunable filter, and then introduced into the sample through the back focal plane of the objective. Using a translation stage, the laser beams were shifted toward the edge of the objective so that emerging light reached the sample at incidence angles slightly smaller than the critical angle of the glass-water interface to illuminate several micrometers into the sample. For STORM imaging, continuous illumination of the 647-nm laser (∼2 kW cm^−2^; for AF647) was used to excite fluorescence from labeled dye molecules and to switch them into the dark state. For PALM imaging, continuous illumination of the 560-nm laser (∼2 kW cm^−2^; for mEos2) was used to excite fluorescence from photoconvertible mEos2 or mMaple3 proteins. Emitted photons were collected from single mEos2 or mMaple3 proteins until they photobleached. Concurrent illumination of the 405-nm laser was used to convert native-state mEos2 or mMaple3 proteins into their photo-converted state. This process results in a cycle of photoconversion and photobleaching for each single protein. The power of the 405-nm laser was adjusted during image acquisition so that at any given instant, only a small, optically resolvable fraction of the proteins in the sample were in the emitting state. For 3D-STORM and 3D-PALM imaging, a cylindrical lens was inserted into the imaging path so that images of single molecules were elongated in *x* and *y* for molecules on the proximal and distal sides of the focal plane (relative to the objective), respectively. STORM and PALM images were taken consecutively, with STORM preceding PALM imaging due to the longer wavelength absorbed by the STORM dye.

#### Image processing

3D STORM and 3D PALM datasets were processed according to previously described methods (3, 6). Briefly, the centroid positions and ellipticities of the single molecule fluorescent spots obtained from raw STORM and PALM data were used to deduce lateral and axial positions of single fluorescent emitters within each sample. Final super-resolution images used in figures result from blurring each single molecule point to a 2D Gaussian profile.

#### Averaging individual synapsed chromosomes

Once 2-color 3D STORM and PALM images of late pachytene nuclei in whole gonads were processed and aligned, individual stretches of synapsed chromosomes from each image were aligned to a consistent orientation for purposes of overlaying and averaging. Individual synapsed axes are then smoothed in *x* and *z* dimensions using a locally weighted scatterplot smoothing algorithm to ultimately produce axes running vertically with no angular deviation in the *xy* plane or *yz* plane. Data from individual synapsed chromosomes is combined and overlaid. By subsequently blurring the data points to a 2D Gaussian profile, images that are effectively averages over several stretches of synapsed chromosome axes are obtained. To determine the localizations of components in the synapsed chromosome axes, histograms in *x* (frontal view) and *z* (cross-sectional view) were fitted to two and one normalized Gaussian distributions, respectively. To compare localizations, we use the positions from the center of the SC in frontal view (*x*) and half widths at half maximum (HWHMs) in vertical view (*z*). Standard deviations of the positions were estimated by bootstrapping, using subsets including half of the individual stretches analyzed per component.

---

**Supplementary Table S1:**
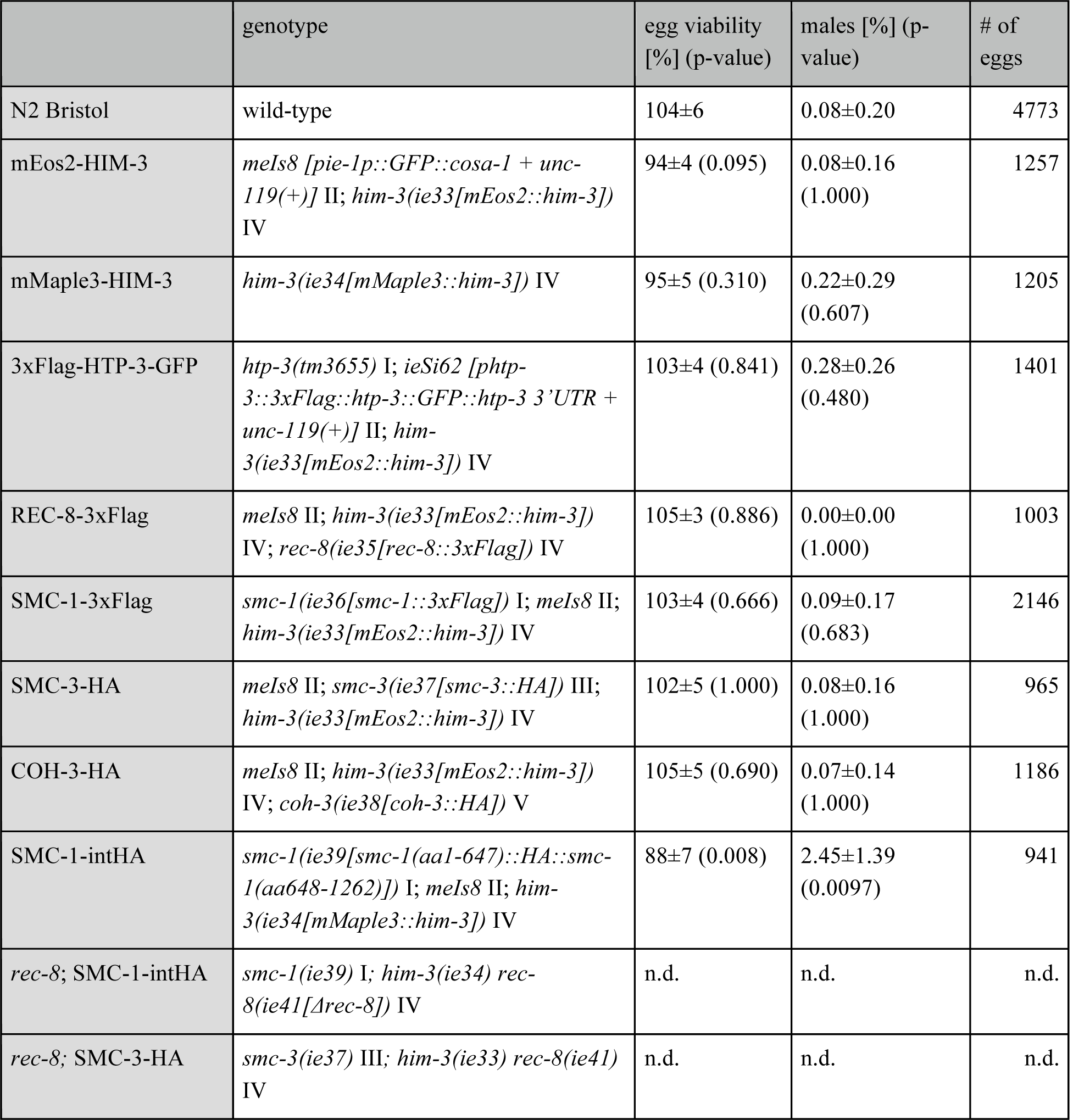
List of strains and egg-counts.

**Supplementary Figure S1:**
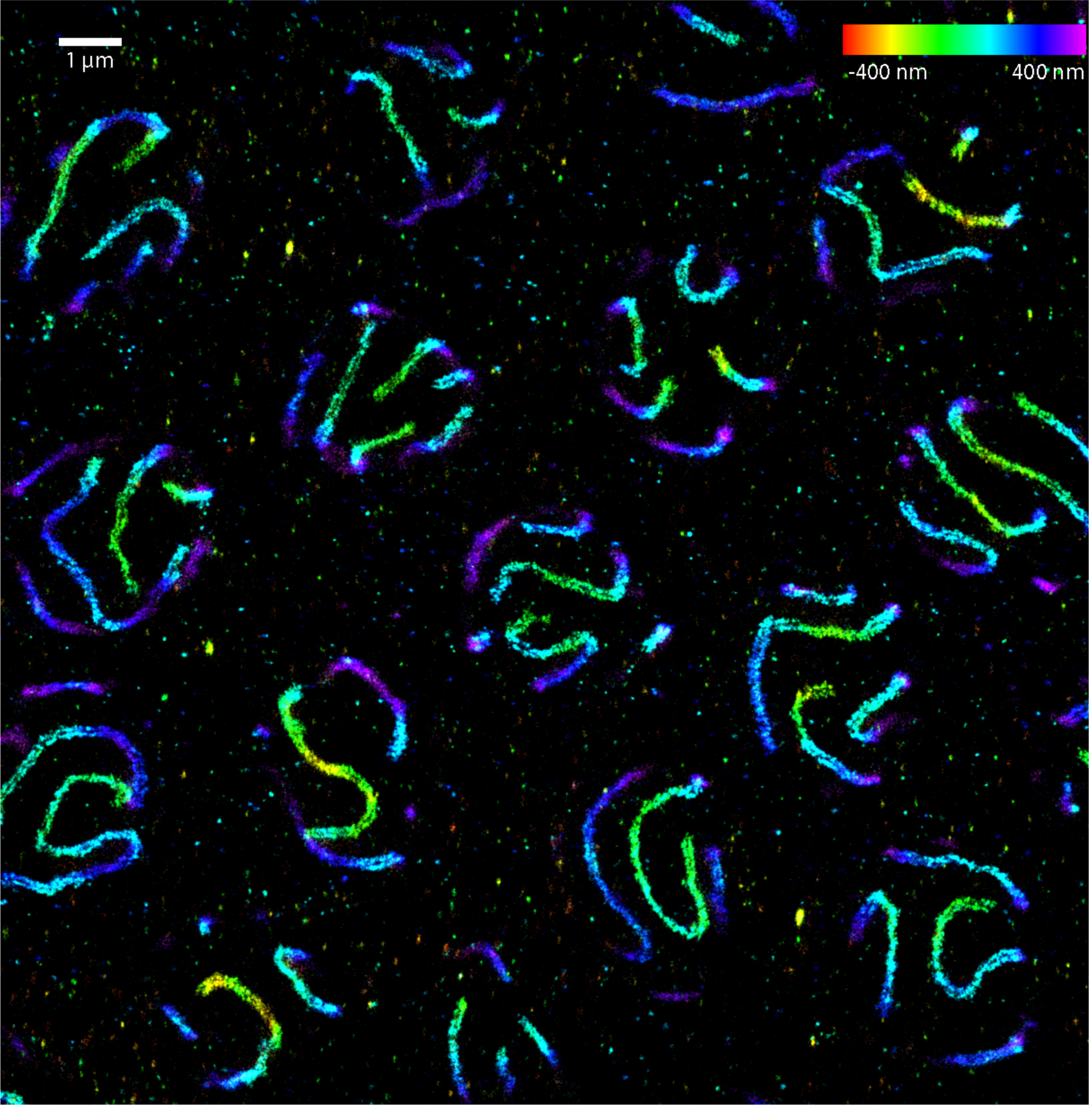
3D-STORM image of HIM-3. Colors denote localization in *z.*

**Supplementary Figure S2:**
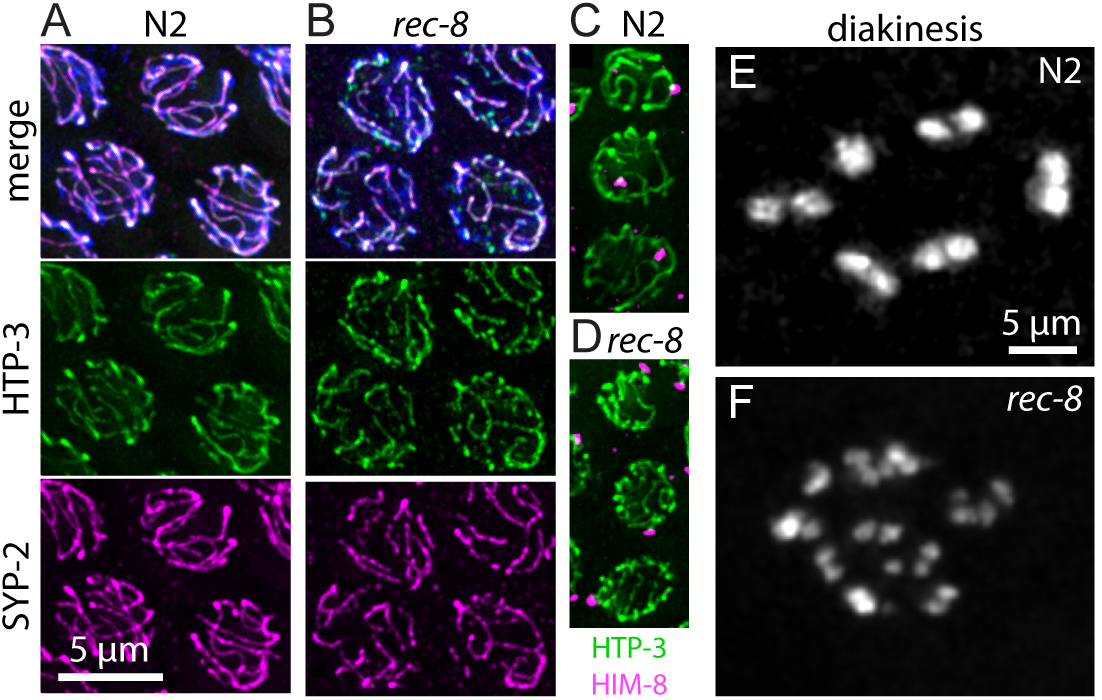
The newly-engineered mutation *rec-8(ie41)* is a null allele of REC-8. Chromosomes undergo synapsis in wild-type (A) and *rec-8(ie41)* mutants (B). Top panel shows a merge of DAPI (blue) and immunostaining of HTP-3 (green, middle panel), SYP-2 (magenta, bottom panel). Synapsis is homologous as demonstrated by the formation of a single HIM-8 focus, which marks chromosome X, in both, wild-type (C) and *rec-8(ie41)* mutants (D). HIM-8 is shown in magenta, HTP-3 in green. In diakinesis, 6 bivalents are observed in wild-type nuclei (E, DAPI staining), while *rec-8(ie41)* nuclei exhibit “butterfly-shaped” univalent chromosomes and occasional fragments (F).

## Supplementary Video S1: Three-dimensional model of the synapsed chromosome axis

A model of the synapsed chromosome axis in three-dimensions was constructed by combining the localizations of domains determined by STORM (position from midline in *x* and HWHM in *z*) and prior information about their interactions (structure of cohesin complexes from (51-54), interactions of HORMA-domain proteins from (20)). The model is rendered using Pov-Ray 3.7.

